# Flexible Protein-Protein Docking with a Multi-Track Iterative Transformer

**DOI:** 10.1101/2023.06.29.547134

**Authors:** Lee-Shin Chu, Jeffrey A. Ruffolo, Ameya Harmalkar, Jeffrey J. Gray

## Abstract

Conventional protein-protein docking algorithms usually rely on heavy candidate sampling and re-ranking, but these steps are time-consuming and hinder applications that require high-throughput complex structure prediction, e.g., structure-based virtual screening. Existing deep learning methods for protein-protein docking, despite being much faster, suffer from low docking success rates. In addition, they simplify the problem to assume no conformational changes within any protein upon binding (rigid docking). This assumption precludes applications when binding-induced conformational changes play a role, such as allosteric inhibition or docking from uncertain unbound model structures. To address these limitations, we present GeoDock, a multi-track iterative transformer network to predict a docked structure from separate docking partners. Unlike deep learning models for protein structure prediction that input multiple sequence alignments (MSAs), GeoDock inputs just the sequences and structures of the docking partners, which suits the tasks when the individual structures are given. GeoDock is flexible at the protein residue level, allowing the prediction of conformational changes upon binding. For a benchmark set of rigid targets, GeoDock obtains a 41% success rate, outperforming all the other tested methods. For a more challenging benchmark set of flexible targets, GeoDock achieves a similar number of top-model successes as the traditional method ClusPro [1], but fewer than ReplicaDock2 [2]. GeoDock attains an average inference speed of under one second on a single GPU, enabling its application in large-scale structure screening. Although binding-induced conformational changes are still a challenge owing to limited training and evaluation data, our architecture sets up the foundation to capture this backbone flexibility. Code and a demonstration Jupyter notebook are available at https://github.com/Graylab/GeoDock.

## 1 Introduction

Protein-protein interactions are involved in nearly all cellular functions in living organisms, from signaling and regulation to recognition. These cellular functions are crucially dependent on the precise assembly of proteins to become functional multi-protein complexes. Understanding the 3D structures of protein-protein complexes at the atomic level can give insight into the mechanisms that underlie these functions. While experimental approaches can determine protein structures, they are expensive, low-throughput, and not applicable to all proteins. Recent breakthroughs of AlphaFold2 [3] and other follow-up works [4–7] have demonstrated the prediction of 3D models of protein structures comparable to experimental accuracy. Along with genome-wide protein sequencing, the AlphaFold database [8] provides open access to 214M protein structure predictions from the sequences deposited in UniProt [9]. To provide proper biological context to the expanding number of predicted protein monomer structures, fast and reliable computational approaches for modeling protein-protein interactions are critical. Protein-protein docking methods provide computational tools for fundamental studies of protein interactions by predicting the favorable protein binding sites and possibly binding-induced conformational changes.

Protein-protein docking methods predict a protein complex structure given the structures of its unbound monomeric partners. Classical protein docking approaches generally comprise (1) a sampling algorithm that generates ensembles of candidate docked structures and (2) a scoring function that evaluates the candidates generated from the sampling stage. The sampling strategies can be classified into exhaustive global search, local shape feature matching, or randomized search [10]. The exhaustive global search methods [11–16, 1, 17] mostly use fast Fourier transforms (FFTs) [18] to cover the complete 6D (3D translational plus 3D rotational) space, assuming no conformational changes of the docking partners. The local shape matching methods [19–24] typically represent a protein by the shape of its molecular surface and find matches of high shape complementarity between two proteins. The randomized search methods [25–28, 2, 29–36] use stochastic algorithms such as Monte Carlo methods to explore docking poses through the free energy landscape starting from a randomized initial pose, with the protein represented as an all-atom or coarse-grain model. Depending on the sampling algorithms used, the scoring can take place after or can be coupled during the sampling process. In contrast to sampling and scoring scheme, template-based docking methods [37] use information such as sequence similarity, evolutionary conservation, and interface complementarity to search for complexes that are homologous to the proteins to be docked, and then use the complexes as docking templates. Due to the massive candidate searching and evaluation, these docking methods are usually time-consuming, hindering applications that require high-throughput complex structure prediction.

Recent breakthroughs in machine learning, especially in deep learning techniques, have been applied to several protein docking-related tasks. MaSIF [38] and ScanNet [39] use geometric deep learning methods to predict the binding sites of a protein based on the structural and chemical features of its molecular surface, independent of the binding partners. For partner-specific protein interface prediction, several methods [40–47] predict protein inter-chain contacts (residues within some cut-off distance) using Graph Neural Networks (GNNs) that input structural and possibly evolutionary features of the proteins. The predicted binding sites or inter-chain contact maps from these deep learning methods can guide docking and generate protein complex structures [48, 49], but the resulting complex structures have not yet achieved accuracy comparable to the conventional docking approaches. End-to-end deep learning methods that predict protein complex structures from protein sequences and multiple sequences alignments (MSAs) have also been developed. These methods [50–52] extend the ability of AlphaFold2 [3] that was originally designed for protein folding to protein multimer structure prediction. Remarkably, AlphaFold-Multimer (AF-M) [53] includes multiple chains during training and outperforms other AlphaFold2-based approaches. However, searching for MSAs slows the inference process, hampering its applications to fast protein complex structure predictions. Furthermore, MSAs do not apply to the docking of important protein families like antibodies or T-cell receptors that evolve at different timescales than their binding partners [54].

Several deep learning methods have been developed for fast rigid docking. Ganea *et al* [55] developed EquiDock, an equivariant graph-matching neural network that predicts the rotation and translation to place one of the proteins at the correct docked position relative to the second protein. Sverrisson *et al* [48] developed a generative model based on the interface features derived from dMaSIF [56] to produce ensembles of docking candidates that are then scored by a trained discriminative model. Due to the rigid-body assumption, these methods are not capable of predicting the conformational changes upon protein binding. In addition, the docking success rates of these methods are lower than the conventional docking methods. More recently, McPartlon & Xu [57] proposed DockGPT for flexible and site-specific protein docking and design, demonstrating high docking success rates when partial binding site information is provided.

In this work, we develop GeoDock, a multi-track iterative transformer network for fast and flexible protein-protein docking. We use the ESM language model [6] to embed the sequences, thereby avoiding MSAs during training. Our approach involves a multi-track architecture inspired by AlphaFold2 [3], and it incorporates the input structure information and the invariant point attention (IPA) module to enable backbone motion. We demonstrate a superior performance of the docking success rate on a benchmark dataset consisting of bound docking tasks. We measure the extent of backbone motion the network can capture, and we highlight limited success on a benchmark set of unbound docking tasks.

## 2 Methods

### Datasets

For training, fine-tuning, and evaluating our models, we used datasets from (1) Database of Interacting Protein Structures (DIPS) [41] and (2) Docking Benchmark 5.5 (DB5.5) [58, 59]. DIPS comprises 42,826 non-redundant experimentally resolved binary protein complexes (< 30% sequence identity to any protein in the DB5.5). We filtered out the complexes with less than 50 residues in a single chain, resulting in train/validation/test splits of size 36318/927/100. DB5.5 contains 271 protein complexes with both bound and unbound conformations. We curated two subsets from DB5.5 for evaluation. The first subset (DB5.5 bound) consists of 24 complexes that were used as the test set in the EquiDock [55] report. The second subset (DB5.5 unbound) comprises 69 complexes that are classified as medium difficulty or difficult targets. We used the remaining 178 complexes for fine-tuning the model pre-trained on DIPS.

### Continuous Cropping for Model Training

We randomly cropped a continuous block of residues on each chain up to 500 residues in total (summing over the chains) to fit the memory size of the GPU (NVIDIA A100 40GB) we used. The cropping strategy ensures that the cropped structure still has two continuous chains, and the length of the chains always sums to 500. Cropping is used for training the models only but not for inference.

### Input Protein Graphs

We represent the input docking partners as two separate graphs. Each graph is fully-connected, with nodes representing the amino-acid identity and edges representing the geometric relations between the intra-chain residues. For node embeddings, we used the ESM-2 (650M) pre-trained language model [6], which inputs a protein sequence and outputs per-residue embeddings. The embeddings from the language model have been shown to improve downstream structure prediction tasks such as protein folding [60, 6]. For edge embeddings, we used the inter-residue distance and orientation distributions adopted from trRosetta [61] and the relative positional encodings from Evans *et al* [53]. The two separate graphs from the two docking partners are concatenated before input to the neural network, with the inter-chain information masked. The graph representation contains rich geometric information without any task-specific or hand-crafted features.

### Inter-chain Contact Information for Model Training

Inspired by the work from McPartlon & Xu [57], we provided a small fraction of inter-chain contact (distance between C_*β*_ atoms under 10 Å) information as a binary embedding *C*_*ij*_ *∈* {0, 1} concatenated to the edge embeddings during training. Specifically, for each training example, we generated a random integer *i ∈* [0, 3] and randomly sampled *i* inter-chain contacts. Thus, during training, the model sees 0 to 3 contacts with equal probability. During inference for the blind docking tests, no inter-chain contact information is provided (*C*_*ij*_ = 0).

### Multi-Track Iterative Transformer Network

We predict a docked protein complex structure from the input docking partners end-to-end via a multi-track iterative transformer network as shown in Figure 1. Inspired by the model architecture of AlphaFold2 [3], the network comprises a graph module followed by a structure module. The graph module communicates 1D (node embeddings) and 2D (edge embeddings) information and outputs the updated embeddings. The structure module inputs the updated embeddings from the graph module along with the backbone frames and outputs the predicted rotations and translations for updating the backbone frames. The main difference between our network and AlphaFold2 is that we input sequences and structures instead of MSAs. In addition, the network is much shallower compared to AlphaFold2, with only 3 layers for both the graph and structure modules. During training, we iterate the network 4 times with no gradient except for the last iteration. During each iteration, we recycle the outputs from both the graph and structure modules. Our implementation of recycling differs from AlphaFold2, as we additionally recycle the inter-residue orientation distribution from the output backbone frames of the structure module. The number of trainable parameters is around 4.3M. The last iteration outputs the predicted backbone frames, pair-wise distance bins, and per-residue confidence scores.

**Figure 1:**
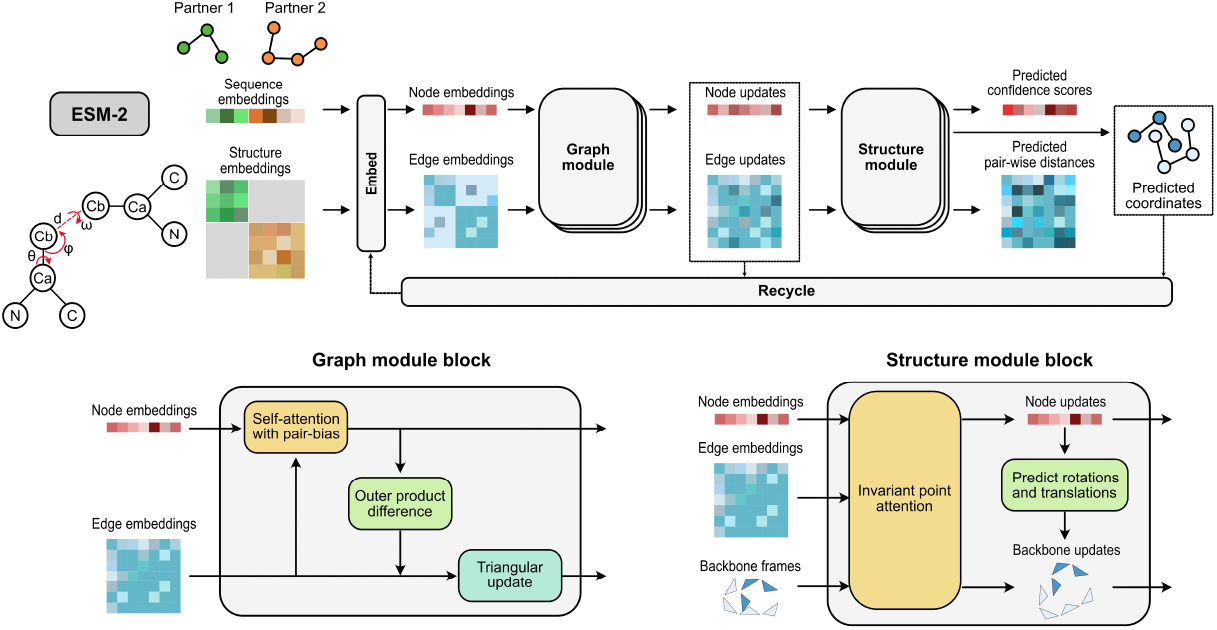
Diagram of the multi-track iterative transformer network for predicting a docked protein complex structure from input docking partners.

### Loss Functions

To train our neural network, we used intra-chain and inter-chain frame-alignment point error (FAPE), pairwise distance loss, the C_*α*_ local distance difference test (lDDT-C_*α*_) [62] loss, and structure violation loss. The loss functions are largely adopted from AlphaFold-Multimer [53], but we omit the cross-entropy loss for sequences, as we do not input MSAs. We also exclude the within-residue violation term in the structure violation loss because our network does not predict side-chain atoms.

### Training Stages

We found that including the inter-chain FAPE at the beginning hinders the convergence of the intra-chain FAPE, as the gradients from the two loss functions compete. Therefore, we trained our network in three stages with gradually increasing loss terms. In the first stage, we only include the intra-chain FAPE, pair-wise distance loss, and lDDT-C_*α*_ loss. In the second stage, we add the inter-chain FAPE. At the last stage, we add the structure violation loss. With this training procedure, the neural network initially learns how to position the intra-chain residues, and subsequently acquires the ability to assemble the docking partners. Our model was first trained on the DIPS training set for 50 epochs, using Adam with a learning rate 5 *×* 10^*−* 5^. The best DIPS-validated model was then fine-tuned on the DB5.5 training set alone with the same optimization settings.

## 3 Results

### Evaluation

We evaluate the predicted structures using the DockQ score [63], which assesses the structural quality of the interface and provides a score ranging from 0 to 1. Following the practice in the CAPRI blind prediction challenge [64], we use a DockQ score threshold of 0.23 to count as a successful dock, corresponding to an “acceptable” CAPRI ranking.

### Bound-conformation Docking Results

For the bound docking task, we compared GeoDock with the existing methods mentioned in EquiDock [55]. Each method belongs to one of the categories specified in the introduction. ClusPro [1] is an exhaustive global search method using FFT; PatchDock [21] is a local shape feature matching approach; ATTRACT [30] is energy minimization with a reduced protein model; EquiDock [55] is a deep learning rigid-body docking approach. The predicted protein complex structures of these methods are taken from the EquiDock GitHub repository https://github.com/octavian-ganea/equidock_public. We excluded HDOCK [65] in our comparison, as it is a template-based docking method and the test targets are in HDOCK’s template set. We cannot compare GeoDock to the dMaSIF rigid docking method [48] since they have not reported the per-target docking success rate, and their code is not accessible. We also do not compare with AlphaFold-Multimer [53] due to the presence of test targets within their training sets. We employed the best DIPS-validated model and the DB5.5 fine-tuned model to evaluate DIPS and DB5.5 held-out test sets, respectively. Our method GeoDock successfully docks 41% of the targets in the DIPS test set, outperforming all the other tested methods (Fig 2A). On the DB5.5 bound set of targets tested in EquiDock [55] paper, GeoDock obtains a 25% success rate, outperforming EquiDock and PatchDock but underperforming other traditional methods (Fig 2B). The lower success rate on DB5.5 relative to DIPS suggests that the DIPS training set, which is predominantly homo-multimers, may not adequately cover the range of interface diversity in the DB5.5 set.

**Figure 2:**
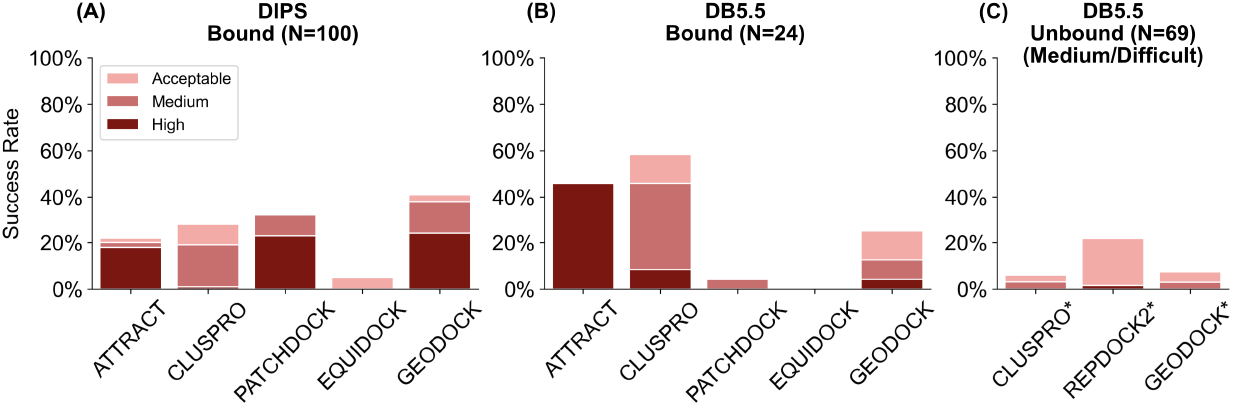
(A) DIPS, (B) DB5.5 bound, and (C) DB5.5 unbound test sets results of docking success rates with CAPRI acceptable, medium, or high quality ranking. The success rates are calculated as the fraction of cases within a specific range of DockQ scores. Scores above 0.23 are considered acceptable, scores between 0.49 and 0.80 are considered medium quality, and scores above 0.80 are considered high quality [63]. * indicates inference starting from the unbound conformation.

### Unbound-conformation Docking Results

For the unbound docking task, we compared GeoDock with the conventional methods ClusPro [1] (rigid docking) and ReplicaDock2 [2] (flexible docking). For ClusPro, we used the single top model success rate reported in their paper [66]. For ReplicaDock2, we selected the single top model (ranked by Rosetta interface score) obtained from blind docking runs. We evaluate our method on the DB5.5 unbound set (Fig 2C) of flexible targets (classified as medium or difficult). The success rate of GeoDock performs similarly to the traditional rigid docking method ClusPro (7% vs 6%), demonstrating GeoDock’s capability of unbound docking. However, GeoDock underperforms the physics-based flexible docking method ReplicaDock2 (7% vs 22%). Similar to GeoDock, DockGPT [57] reported a blind unbound docking success rate (7%) on a DB5.5 set of 42 targets (classified as rigid, medium, or difficult). The generally lower docking success rates for unbound docking reflect the current difficulty of deep-learning docking approaches to capture backbone conformational changes.

### Backbone Flexibility

As shown in Figure 3, GeoDock is able to move the backbone at the flexible regions. The current GeoDock model typically pushes the backbone away from the bound state (Fig 3B) similar to previous conformer selection methods [67]. The observed movement is minimal (< 1Å Fig 3A) and the predicted structures closely resemble the input unbound structures (Fig 3C). The current training examples had no conformational change, so a next step might be to fine-tune our model with an ensemble of sampled backbone conformations (e.g. from ReplicaDock2 [2]) to better capture conformational changes from the unbound to bound states.

**Figure 3:**
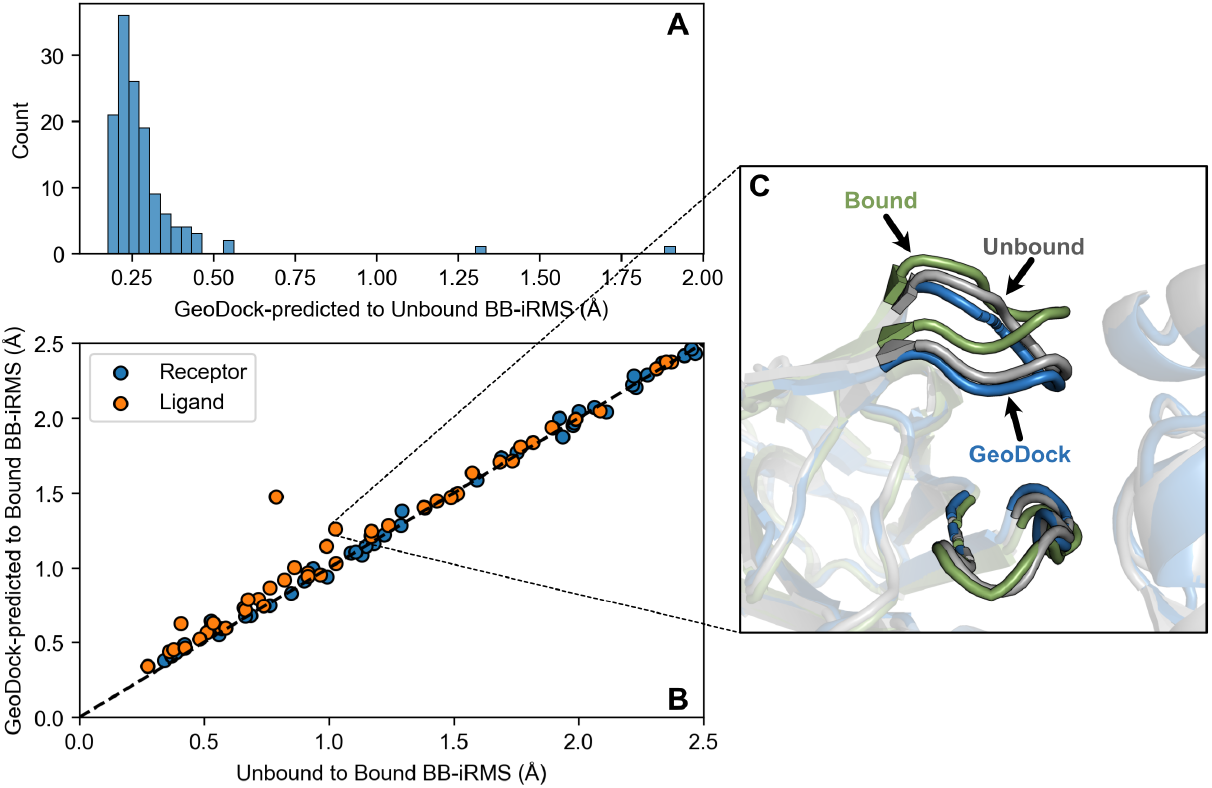
Analysis of backbone flexibility measured with backbone interface RMSD (BB-iRMS) for DB5.5 flexible targets. (A) BB-iRMS distributions of GeoDock-predicted structures to unbound structures. (B) BB-iRMS of GeoDock-predicted structures to bound structures versus unbound to bound. (C) Comparison of the unbound structure (grey) superimposed over the bound (green), and the GeoDock-predicted structure (blue) for the DsbC-DsbDalpha complex (PDB accession code: 1JZD_AB:C (bound); 1JZO_AB and 1JPE_A (unbound)).

### Inference Speed

The inference times of GeoDock are 0.76 *±* 0.62 seconds for the DIPS test set (residue sizes 100-800) and 0.80 *±* 0.52 seconds for the DB5.5 test set (residue sizes 200-700) on a single NVIDIA V100 GPU, which is faster than the reported runtimes for EquiDock (5 *±* 5 and 5 *±* 10 seconds) and is at least 10^3^ times faster than the other methods [55]. Inference time is important for applications that require high throughput protein complex structure prediction, such as structure-based virtual screening.

## 4 Discussion and Conclusion

GeoDock is a fast, end-to-end approach for flexible protein-protein docking with a multi-track iterative transformer network. Unlike previous deep learning methods for protein complex structure prediction that require MSAs as input [3, 4], GeoDock inputs structure and sequence information of individual proteins and predicts a docked protein complex structure, which suits the task when the unbound structures are given. Furthermore, the model allows for backbone movements, which is in contrast to previous deep learning methods that assume no conformational changes [55, 48]. With the same training dataset and faster inference speed, our method GeoDock outperforms EquiDock [55] on both DIPS and DB5.5 test sets. The observed improvement can be attributed to the iterative updating of coordinates, which contrasts with the EquiDock paper’s approach of predicting a single transformation for one of the docking partners.

While this method significantly accelerates the inference process, in terms of docking success rates, it does not surpass conventional methods that use sampling and re-ranking strategies. This may be attributed to the training objective of the model not aligning with the evaluation of docking success, which is determined by the percentage of predictions with a DockQ score above a certain threshold. The objective function of GeoDock is to minimize the expected error between the predicted and ground truth structures. Trained with this objective, the model learns to predict structures with low average error, although each individual structure may not necessarily be considered a success case if the error is not sufficiently low. A more viable objective is to identify a low-error docking pose distribution and sample from it, which can be framed as a generative model similar to a prior study on the small molecule docking using diffusion model [68]. This suggests a future research direction of deep generative models in sampling protein docking poses.

We have demonstrated that GeoDock can move the backbone despite being currently trained only on the bound protein complexes. However, the movement is minimal, and the predicted structures closely resemble the input unbound structures. We leave for future work the fine-tuning of the model with sampled backbone conformations to test whether the model can better capture the conformational change upon binding. This is particularly important for cases such as allosteric inhibition or docking from uncertain unbound model structures, where the flexible regions on the proteins hinder the success of rigid docking.

For target-specific protein binder design, e.g., screening antibodies for a specific antigen, a high-throughput docking algorithm is usually required to scan over a vast search space of potential hits. With further development, GeoDock can serve as a fast and flexible protein-protein docking tool and facilitate the design of protein binders and drugs for a wide variety of targets.

## Acknowledgements

This work was supported by National Institutes of Health grants R35-GM141881. Computational resources were provided by the Advanced Research Computing at Hopkins (ARCH).

## Code Availability

Code, pre-trained models, and a demonstration Jupyter notebook for GeoDock are available at https://github.com/Graylab/GeoDock.

## Notes

### Competing Interest Statement

Jeffrey Gray is an unpaid board member (co-director) of the Rosetta Commons. Under institutional participation agreements between the University of Washington, acting on behalf of the Rosetta Commons, Johns Hopkins University may be entitled to a portion of revenue received on licensing Rosetta software including some methods described in this study.
Jeffrey Gray has a financial interest in Cyrus Biotechnology. Cyrus Biotechnology distributes the Rosetta software, which may include methods described in this study. The results of the study discussed in this article could affect the value of Cyrus Biotechnology.
These arrangements have been reviewed and approved by the Johns Hopkins University in accordance with its conflict-of-interest policies.

